# Fast Nonnegative Matrix Factorization and Applications to Pattern Extraction, Deconvolution and Imputation

**DOI:** 10.1101/321802

**Authors:** Xihui Lin, Paul C. Boutros

## Abstract

Nonnegative matrix factorization (NMF) is a technique widely used in various fields, including artificial intelligence (AI), signal processing and bioinformatics. However existing algorithms and R packages cannot be applied to large matrices due to their slow convergence, and cannot handle missing values. In addition, most NMF research focuses only on blind decompositions: decomposition without utilizing prior knowledge. We adapt the idea of sequential coordinate-wise descent to NMF to increase the convergence rate. Our NMF algorithm thus handles missing values naturally and integrates prior knowledge to guide NMF towards a more meaningful decomposition. To support its use, we describe a novel imputation-based method to determine the rank of decomposition. All our algorithms are implemented in the R package NNLM, which is freely available on CRAN.

## 1 Introduction

Nonnegative matrix factorization (NMF or NNMF) ([Lee and Seung, 1999]) has been widely used as a general method for dimensional reduction and feature extraction on nonnegative data. The main difference between NMF and other factorization methods, such as SVD, is the nonnegativity, which allows only additive combinations of intrinsic ‘parts’, *i.e.* the hidden features. This is demonstrated in ([Lee and Seung, 1999]), where NMF learns parts of faces and a face is naturally represented as a additive linear combination of different parts. Indeed, negative combinations are not as intuitive or natural as positive combinations.

In bioinformatics, NMF is sometimes used to find ‘meta-genes’ from expression profiles, which may be related to some biological pathways ([Brunet *et al*., 2007], [Kim and Park, 2007]). NMF has been used to extract trinucleotide mutational signatures from mutations found in cancer genomic sequences and it was suggested that the trinucleotide profile of each cancer type is positive linear combination of these signatures ([Alexandrov *et al*., 2013]).

There are several different algorithms available for NMF decomposition, including the multiplicative algorithms proposed in ([Lee and Seung, 1999]), gradient descent and alternating nonnegative least square (ANLS). ANLS is gaining attention due to its guarantee to converge to a stationary point and faster algorithms for nonnegative least squares (NNLS).

In this paper, we adapt the ANLS approach, but incorporate a solution to the NNLS problem using a coordinate-wise algorithm proposed by [Franc *et al*., 2005], in which each unknown variable can be solved sequentially and explicitly as simple quadratic optimization problems. We demonstrate that this algorithm can converge much faster than traditional multiplicative algorithms. For NMF with Kullback-Leibler divergence loss, we extend this methodology by approaching the loss with a quadratic function.

NMF is a dimension reduction method, as the resulting decomposed matrices have a smaller number of entries than the original matrix. This means that one does not need all the entries of original matrix to perform a decomposition, which means that NMF should be able to handle missing values. Furthermore, the reconstructed matrix have values on the entries where the original matrix has missing values. This reveals the capability of NMF for missing value imputation. Inspired by this capability and the popular training-validation tuning strategy in supervised models, we introduce a novel method to optimize the only hyper-parameter *k*, *i.e.* the rank of NMF decomposition.

NMF is essentially unsupervised. It performs a blind decomposition, which puts the meaning of the result in question. This can limit the applications of unsupervised methods in areas in where strong interpretability is critical, including most biomedical research. On the other hand, decomposition without utilizing known discovery (prior knowledge) may not be effective, especially with a small sample size. To overcome these challenges, we apply a mask technique to the NMF decomposition during the iterating algorithms to retain certain structures or patterns in one or both resulting matrices, which can be designed according to our prior knowledge or research interest. This technique can be used to perform a pathway or subnetwork guided decomposition, or to separate different cell types from mixed tissue samples.

All of these algorithmic innovations are implemented in the popular R programming language. They serve as an alternative to the widely-used *NMF* package ([Gaujoux and Seoighe, 2010]) which was first translated from a MATLAB package and later optimized via C++ for some algorithms. The sparse alternating NNLS (ANLS) by ([Kim and Park, 2007]) is anticipated to be algorithmically fast, but is implemented in R leading to slow performance in practice. Our NNLM package combines the efficient NNLS algorithm with the use of Rcpp, which seamlessly integrates R and C++ ([Eddelbuettel and Francois, 2011]) and is freely available and open-source.

## 2 Approach

A typical nonnegative matrix factorization problem can be expressed as 
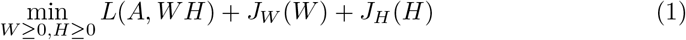
where 
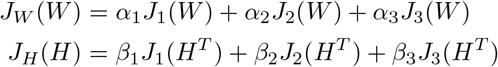
and 
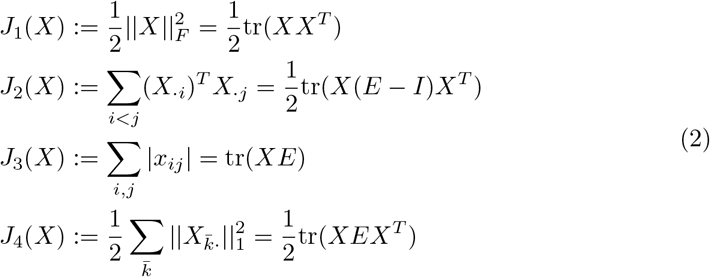

Here *A* ∊ ℝ^*n*×*m*^, *W* ∊ ℝ^*n*×*K*^, *H* ∊ ℝ^*K*×*m*^, *I* is an identity matrix, *E* is a matrix of proper dimension with all entries equal to 1, *X_·i_* and *X_i·_* are the *i*^th^ column and row respectively. Obviously, *J*_4_ = *J*_1_ + *J*_2_. *L*(*x, y*) is a loss function, which is mostly chosen to be square error 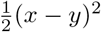, or KL divergence distance *x* log(*x*/*y*) − *x* + *y*. The latter can be interpreted as the deviance from a Poisson model.

The above four types of regularizations can be used for different purposes. *J*_1_ is a ridge penalty to control the magnitudes and smoothness. *J*_2_(*X*) is used to minimize correlations among columns, *i.e.*, to maximize independence or the angle between *X_·i_*, *X_·j_* ([Zhange *et al*., 2008]). *J*_3_ and *J*_4_ ([Kim and Park, 2007]) is a LASSO-like penalty, which controls both magnitude and sparsity. *J*_3_(*X*) tends to control matrix-wise sparsity, which may result in some columns of *X* having all entries equal to 0, while *J*_4_(*X*) forces sparsity in a column-wise manner.

An alternating algorithm solves *W* and *H* alternately and iteratively. Due to the nonnegative constraint, the penalty does not bring additional complexity. In addition, convergence is guaranteed as the loss is monotonically non-increasing at each step. It converges to a local minimum, as if not, there exists a coordinate with non-zero derivative and thus the correspondent update will have make an improvement on the loss.

## 3 Methods

In this section, we will introduce a new and fast algorithm for NMF with mean square loss (section 3.1) and KL-divergence distance loss (section 3.2) with regularization, based on the alternating scheme. In section 3.3, we generalize the multiplicative algorithm proposed by ([Lee and Seung, 1999]) to incorporate all the regularizations in Equation (2). We then re-examine NMF in section 3.4 and develop a novel method to integrate prior knowledge into NMF and guide the decomposition in a more biologically meaningful way, which can be powerful in applications. In section 3.5, we show how one can utilize NMF to impute missing values in array data, and such a capacity leads to a very intuitive and logically robust tuning method to determine the unknown parameter *k* (the rank) in section 3.6.

### 3.1 Alternating nonnegative least square (ANLS)

When *L* is a square loss, the following sequential coordinate-wise descent (SCD) algorithm is used to solve a penalized NNLS for *H* while *W* is fixed.

> 0. Let *V* = *W^T^W* + *β*_1_*I* + *β*_2_(*E* − *I*).
>
> 1. Initialization. Set *H*^(0)^ = 0, *U*^(0)^ = −*W^T^ A* + *β*_3_*E*.
>
> 2. Repeat until convergence: for *j* = 1 to *m*, *k̄* = 1 to *k*

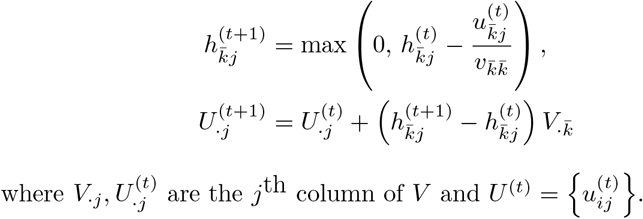

Since *E − I* is semi-negative definite, to ensure the uniqueness and the convergence of the algorithm, we impose the constraint that *β*_1_ ≥ *β*_2_.

The alternating algorithm fixes *W* and solve for *H* using NNLS, and then fixes *H* and solve for *W* using the same algorithm. This procedure is repeated until the change of *A − W H* is sufficiently small.

Instead of initializing *H*^(0)^ = 0 for every iteration, we use a *warm-start*, *i.e.*, make use of the result from previous iteration,

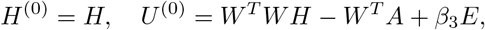
where *H* is the current matrix during the iteration.

### 3.2 Sequential quadratic approximation for Kullback-Leibler divergence loss

When *L* is a KL divergence distance, we use a similar SCD algorithm, by approximating KL(*A*|*W H*) with a quadratic function.

Assume *W* is known and *H* is to be solved. Let 
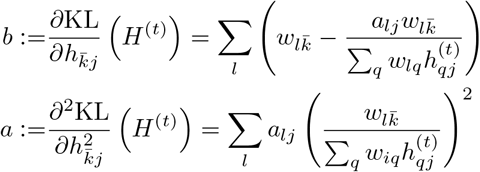
where *H*^(*t*)^ is the current value of *H* in the iterative procedure.

When expanding the penalized KL divergence up to the 2nd order with respect to 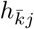 around its current value 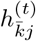 while fixing all others, we can solve the resulting quadrature explicitly by 
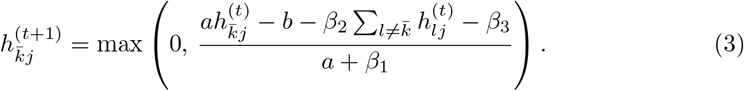

A similar formula for updating 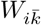 can be derived. Note that when an entry of *Â* is 0, the KL divergence is infinity. To avoid this, we add a very small number both to *A* and *Â*.

### 3.3 Adding regularization to Lee’s multiplicative algorithm

Two multiplicative updating algorithms are proposed in [Lee and Seung, 1999] for square loss and KL divergence loss. We modify these algorithms to integrate all the above regularizations as follows.

With square loss 
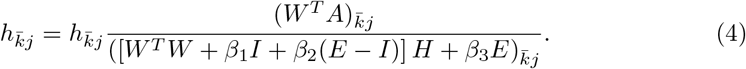

With Kullback-Leibler divergence distance, 
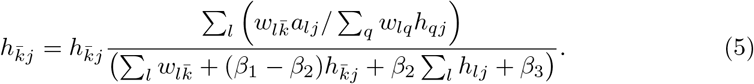

When*β_i_* = 0, *i* = 1, 2, 3, these are the original multiplicative algorithms in [Lee and Seung, 1999]. The updating rules for *W* are similar to (4) and (5).

This multiplicative algorithm is straight-forward to implement, but it has the drawback that when *W* or *H* is initialized with a zero or positive entry, it remains zero or positive throughout the iterations. Therefore, true sparsity can not be achieved generally, unless a hard-thresholding is imposed, as many of entries would be small enough to be thresholded to zero.

### 3.4 Content deconvolution and designable factorization

Microarrays are popular techniques for measuring mRNA expression. Strictly speaking, an mRNA profile of a certain tumour sample is typically a mixture of cancerous and healthy profiles as the collected tissues are ‘contaminated’ by healthy cells. A pure cancer profile is usually more suitable for down stream analysis ([Quon *et al*., 2013]).

One can utilize NMF for such a purpose by formatting it as 
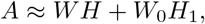
where *W* is an unknown cancer profile, and *W*_0_ is a known healthy profile. Rows of *A* represent probes or genes while columns represent patients or samples. The task here is to solve *W*, *H* and *H*_1_ given *A* and *W*_0_, which can be thought of as a ‘guided’ NMF. In this decomposition, the number of columns of *W* can be interpreted as the number of unknown cancerous profiles or cell types. The corresponding tumour percentage of sample *j* can be estimated as 
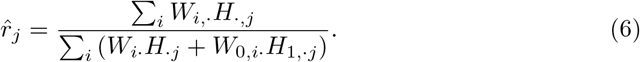

A more general implementation is to use mask matrices for *W* and *H*, where the masked entries are fixed to their initial values or 0 if not initialized. Indeed, one can treat this as a form of hard regularization. It can be seen immediately that the above deconvolution is a special case of this mask technique, in which the masked entries are initialized to the known profile and fixed. This feature is designed to integrate domain knowledge, such as gene subnetworks, pathways etc, to guide the NMF towards a more biologically meaningful decomposition.

For example, assume 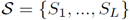, where each *S_l_, l* = 1, …, *L* is a set of genes in certain subnetwork or pathway *l*. One can design *W* as a matrix of *K* columns (*K ≥ L*), with *w_il_* = 0 when *i* ∉ *S_l_*. NMF factorization will learn the *weight* or *contribution w_il_* of real gene *i* in sub-network or pathway *l* from data. One can also interpret *h_lj_* (∑_*i*_ *w_il_*) as an expression level of sub-network or pathway *l* in patient *j*. Beside, *W_·j_*’s for *j* = *L* + 1, …, *K* are unknown sub-networks or pathways. Note that *K* is unknown before hand, but later we will see that it can be determined by NMF imputation.

Similarly, if *S_l_* is a set of marker genes (those that are known to be expressed only in a specific cell type / tissue), for tissue *l* and by letting *w_il_* = 0, 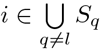, one can find the relative abundance of each tissue type in a sample. A similar formula to Equation (6) can be used to compute the proportions of known (*j* = 1, …, *L*) and unknown (*j* = *L* + 1, …, *K*) cell types / tissues.

Another possible application is for meta-analysis of different cancer types, to find meta-genes that are shared among cancers. For instance, assume *A*_1_, *A*_2_ are expressions of lung cancer and prostate cancer microarrays. By setting certain parts of the coefficient matrix *H* to 0, for example, 
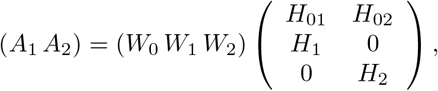
we can expect that *W*_1_ and *W*_2_ are lung and prostate cancer specific profiles, while *W*_0_ is a shared profile.

**Table 1.** A comparison of different imputation methods.

|  | Baseline | Medians | MICE | MissForest | NMF |
| --- | --- | --- | --- | --- | --- |
| MSE | 4.4272 | 0.5229 | 0.9950 | 0.4175 | 0.4191 |
| MKL | 0.3166 | 0.0389 | 0.0688 | 0.0298 | 0.0301 |
| Time (Sec.) | 0.0000 | 0.0000 | 90.2670 | 42.4010 | 0.1400 |

Imputations on a subset of NSCLC microarray data, which composes 200 genes and 100 samples. 30% of the entries are randomly deleted, *i.e.*, missed. MSE = mean square error, MKL = Mean KL-divergence distance and Time = user time.

**Table 2.** Compare performance of different algorithms on a subset of a non-small cell lung cancer dataset, with *k* = 15.

|  | SCD-MSE | LEE-MSE | LEE-MSE-1 | SCD-MKL | LEE-MKL |
| --- | --- | --- | --- | --- | --- |
| MSE | 0.155 | 0.1565 | 0.1557 | 0.1574 | 0.1579 |
| MKL | 0.01141 | 0.01149 | 0.01145 | 0.01119 | 0.01122 |
| Rel. tol. | 1.325e-05 | 0.0001381 | 0.000129 | 6.452e-08 | 9.739e-05 |
| Total epochs | 5000 | 5000 | 5000 | 5000 | 5000 |
| Time (Sec.) | 1.305 | 1.35 | 8.456 | 49.17 | 41.11 |

MSE = mean square error; MKL = mean KL divergence; Rel. tol. = relative tolerance. Elapsed time = actual running time. SCD-MSE = SCD algorithm with MSE loss and 50 inner iterations and LEE-MSE-1 = Lee’s algorithm with MSE loss and 1 inner iteration, *i.e.*, the original multiplicative algorithm.

### 3.5 Missing value imputation

Since matrix *A* is assumed to have a low rank *k*, information in *A* is redundant for such a decomposition. Hence it is possible to allow some entries in *A* to be missing. Such an observation reveals the capability of NMF imputing the missing entries in *A*, which is a typical task in recommendation systems.

The advantage of NMF imputation over traditional statistical imputation methods is that it takes into account all the complete entries when imputing a single missing entry, which implies that NMF can capture complex dependency among entries, while a typical missing value imputation algorithm usually models missing entries in a feature-by-feature (column-by-column or row-by-row) manner and iterates over all features multiple times to capture complex dependency. A comparison of different imputation methods are shown in Figure 1 (B). A subset of NSCLC dataset is used with 30% set to missing at random. One can see that NMF is almost as good as missForest ([Stekhoven and Buehlmann, 2012]) but much faster, and clearly better than MICE and simple median imputation in this example.

**Figure 1.**
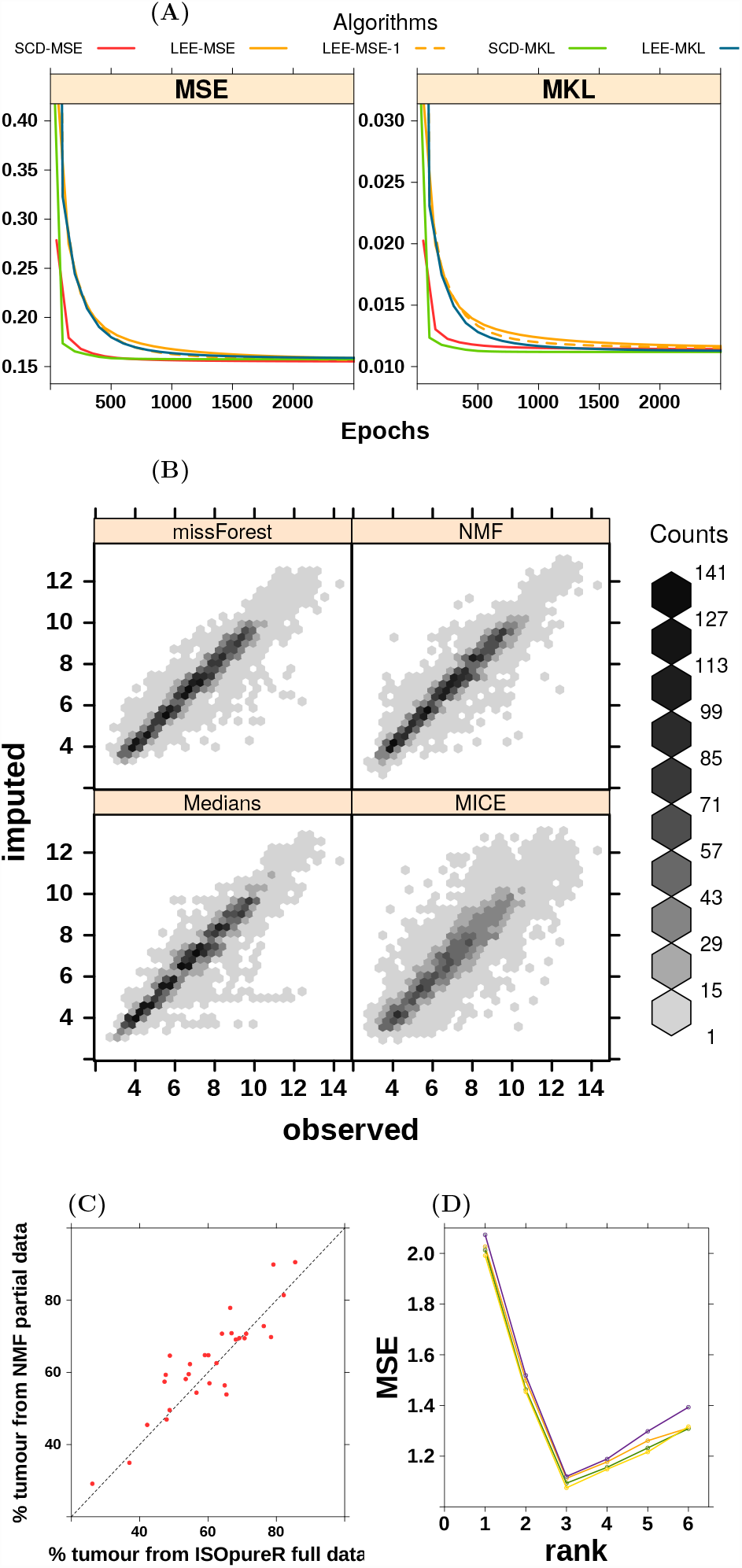
(**A**): Comparison of different algorithms in convergence. (**B**): Comparison of imputation methods. *k* = 2 is used for NMF. (**C**): Compare NMF to ISOpure ([Anghel *et al*., 2015]) for tumour content deconvolution. (**D**): Determine optimal rank *k* in NMF using imputation.

### 3.6 Determine rank via missing value imputation

Tuning hyper-parameter is a typical challenge for all unsupervised learning algorithms. The rank *k* is the only but very crucial parameter, which is *a priori* unknown. [Brunet *et al*., 2007] suggests to try multiple runs of each *k* and uses a consensus matrix to determine *k*. This idea assumes that cluster assignment is stable from run to run if a clustering into *k* classes is strong. However this assumption needs to be validated and the purpose of NMF is not always for clustering. Another idea, brought from the denoising auto-encoder ([Vincent *et al*., 2008]), is to add noise to the matrix *A*, factorize the noisy version and compare the reconstructed matrix to the original *A*. One can expect that the desired *k* should give the smallest error rate. This could be a general approach for many unsupervised learning algorithms. However, when comes to NMF, the choice of ‘noise’ is not easy since the noisy version of *A* has to be nonnegative as well, which suggests that ‘noise’ may also introduce bias.

Given the powerful missing value imputation in NMF, we come up with a novel approach, akin to the well known idea of training-validation split idea in supervised learning. Some entries are randomly deleted from *A* and then imputed by NMF with a set of *k*’s. These imputed entries are later compared to their observed values, and the *k* that gives the smallest error will be our choice, as only the correct *k*, if exists, has the right decomposition that can recover the missing entries. In contrast to the training-validation split in supervised learning, due to the typically big number of entries in *A*, we generally have a very large ‘sample size’. One can also easily adapt the idea of cross-validation to this approach. This idea should be applicable to any unsupervised learning methods that support missing value imputation.

To illustrate, we performed a simulation study. *W* and *H* with *k* = 3, were generated and *A* was constructed by *W H* plus some noise. This result is shown in Figure 1 (D). As we could see, different runs give consistent results. The mean square errors(MSEs) decrease as rank *k* increase, but the increases rate slows down when *k* = 3 (the true rank). Meanwhile, the MSEs for imputed values are minimized at *k* = 3 for all runs.

## 4 Discussion

Both SCD and Lee’s multiplicative algorithms with square error loss have complexity of 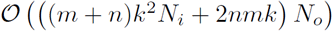, while their KL counterparts have complexity of 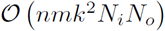, where *N_i_* is the number of inner iterations to solve the nonnegative linear model and *N_o_* is the number of outer iterations to alternate *W* and *H*. *N_i_N_o_* is the total number of epochs, *i.e.*, one complete scan over all entries of *W* and *H*. Obviously algorithms with square error loss are faster than the KL based ones (by a factor of *k*) in terms of complexity, and can benefit from multiple inner iterations *N_i_* (reducing the expensive computation of *W^T^ A* and *AH^T^*) as typically *k ≪ m, n*, which generally should reduce *N_o_*. In contrast, algorithms with KL loss cannot benefit from inner iterations due to the re-calculation of *W H* on each inner iteration. Though the SCD and Lee’s algorithm are similar in terms of complexity, one can expect a much faster convergence in SCD. This is because Lee’s algorithm is essentially a gradient descent with a special step size ([Lee and Seung, 1999]) which is a first order method, while SCD is a Newton-Raphson like second order approach.

Expression deconvolution is of constant interest in bioinformatics and clinical research ([Zhange *et al*., 2008], [Abbas *et al*., 2009]). Some NMF related methods were proposed ([Gaujoux and Seoighe, 2011]). However, our unique methods of using mask matrices are more flexible and powerful, as one can guide the decomposition towards almost arbitrary biological procedure of interest by integrating prior knowledge into the initial and mask matrices. As compared to Bayesian methods like the ISOpure ([Quon *et al*., 2013]), NMF based methods are much faster. An application to an NSCLC dataset was performed and compared to the existing method ISOpure ([Anghel *et al*., 2015]) in Figure 1 (C).

All methodologies described in this paper are implemented in the R package NNLM, available on CRAN.

## 5 Conclusion

We extended the sequential coordinate-wise descent (SCD) to KL-divergence and applied SCD to NMF based on the alternating scheme. We introduced mask matrices in NMF algorithms to integrate prior knowledge to the decomposition, in order to induce more meaningful results. Due to the dimension reduction nature, we found that NMF is an outstanding method for missing value imputation, allowing us to introduce a novel method for tuning hyper-parameters, which is also applicable to any other unsupervised method that can handle missing values.

## Acknowledgements

The authors thank all members of the Boutros lab for supports, especially Dr. Kenneth Chu and Dr. Catalina Anghel.

